# Breeding males, but not females, have elevated androgen receptor expression in the northern house wren (*Troglodytes aedon*), a temperate songbird with female song

**DOI:** 10.1101/2025.09.05.674472

**Authors:** John Yoo, Douglas Weiser, Timothy A. Liddle, Elizabeth M. George, Chelsea Haakenson, Nora H. Prior, Cara A. Krieg, Gregory F. Ball, Karan J. Odom

## Abstract

In many male temperate-breeding songbirds, increased plasma testosterone in the early breeding season regenerates song control nuclei that regulate song. Females of some temperate species also sing but have lower circulating testosterone concentrations. We hypothesized that upregulation of steroid receptors in females could compensate for low circulating testosterone, focusing on the northern house wren (*Troglodytes aedon*), a temperate-breeding songbird in which both sexes sing. We collected brain tissue from both sexes during the early breeding, late breeding, and nonbreeding season. Using quantitative PCR, we quantified mRNA expression of four genes—androgen receptor (AR), estrogen receptors (ERα and ERβ), and aromatase (AROM)—in three song nuclei—HVC, Area X, and RA—and compared sexes. We found females had lower expression than males of AR, AROM and ERα in most song nuclei, especially of AR in HVC in the breeding season. Both sexes, however, had low ERα expression in Area X and RA. In males, expression differed seasonally: breeding males had higher expression of AR in RA and AROM in HVC than nonbreeding males. In both sexes, expression differed among song nuclei: in most genes, HVC had the highest expression, followed by RA, then Area X. These findings suggest singing female house wrens do not compensate for low plasma testosterone by upregulating steroid receptors beyond male levels or during early breeding, when they sing most. Conversely, increased AR expression in breeding males indicates differences in the mechanisms regulating male and female song, with testosterone playing a greater role in male birdsong.

## INTRODUCTION

Testosterone plays a major role in regulating singing of many male northern temperate-breeding songbirds (Ball and Balthazart, 2020; Catchpole and Slater, 2008). In male birds, the traditional view is that increasing photoperiod promotes gonadal growth and secretion of testosterone into plasma (Ball et al., 2004; Bernard et al., 1997). Elevated plasma testosterone, in turn, stimulates growth of the song control system—a network of interconnected brain nuclei that regulate birdsong—through neurogenesis (Nottebohm, 1981; Smith et al., 1997). Key song control nuclei in the song control system, including HVC (acronym serves as proper name), Area X of the medial striatum, and RA (robust nucleus of the arcopallium), express androgen and estrogen receptors, allowing testosterone and its metabolites to regulate gene expression in these regions (Gahr, 2014; Gahr et al., 1993; Nottebohm et al., 1976, 1982; Rose et al., 2022). Testosterone has pleiotropic effects across these brain regions - regulating different components of birdsong, including vocal learning (Area X and HVC), song complexity (HVC), song stereotypy (RA), and song production (RA; Arnold, 1975; Ball et al., 2004; Balthazart and Ball, 2016; Bottjer and Hewer, 1992; Rasika et al., 1994; Smith et al., 1997). Therefore, testosterone can exert multifaceted effects on birdsong (Alward et al., 2017; Ball et al., 2004). This has made male songbirds a successful model system for understanding the neuroendocrine regulation of vocal behavior and learning (Ball and Balthazart, 2020).

In contrast, female song and its neuroendocrine mechanisms have been understudied compared to male song (Riebel et al., 2019; Rose et al., 2022; Rouse, 2022). Historically, researchers viewed song as sexually selected in males, but in many modern songbird species, females sing (Odom and Benedict, 2018; Odom et al., 2025). Moreover, females sang in the common ancestor of all songbirds (Odom et al., 2014). Female song was likely missed in part because female temperate-breeding songbirds often sing at reduced rates (Moyer et al., 2022), for shorter periods of the year (Krieg and Getty, 2016; Wilkins et al., 2020), or not at all (Odom et al., 2014). Consistent with these behavioral sex differences, temperate female songbird song control nuclei are generally less developed in measures such as volume, number of neurons, and neuron size (Krieg and Wade, 2023). In addition, temperate breeding female songbirds often have lower plasma testosterone (Ketterson et al., 2005). The fact that females sing despite these sex differences in neuroanatomy and plasma testosterone suggests that studying the regulation of natural female song may answer questions that studying males alone cannot, including sensitivity of the avian song control system to testosterone. One possible mechanism to explain natural female song in the absence of high circulating plasma testosterone is that females could have increased tissue-specific density of steroid receptors, thus increasing sensitivity to steroid hormones (Gahr, 2014; Gahr et al., 2008; Rosvall, 2013; Rosvall et al., 2012).

The extent to which female song depends on testosterone is unclear. On one hand, testosterone may have similar effects in both sexes. In tropical species, where female song is more common, measures of the song control regions are less dimorphic than in temperate species (Rose et al., 2024), though male song control nuclei are, in general, still larger than those of females (Ball and Macdougall-Shackleton, 2001; Brenowitz and Arnold, 1985; Gahr et al., 2008). In addition, implanting testosterone in female birds increases volumes of song control nuclei and results in development of male-like song (Glisson et al. *In prep*; Madison et al., 2015; Nottebohm, 1980). These studies suggest females are sensitive to testosterone but are still able to sing with lower concentrations of plasma testosterone and less-developed song control regions (e.g., Krieg et al. *in review*; Krieg and Wade 2023). However, these studies do not confirm that females use testosterone-dependent mechanisms to achieve song. Lab studies involving implantation of testosterone in females often administer much higher concentrations of testosterone than females naturally achieve in wild populations (Goymann and Wingfield, 2014). Even in males, testosterone’s effect can be emulated with other stimuli such as photoperiod or social environment (Alward et al., 2014; Ball et al., 2004), and two separate studies tracking neurogenesis in the song control system of two different wild songbird species found that the song control system reaches nearly full volume prior to pre-breeding peaks in circulating testosterone (Caro et al., 2005; Tramontin et al., 2001). This suggests that, even in males, other factors beyond circulating testosterone drive changes in song control nuclei volume. Finally, as most research has been conducted in zebra finches and canaries, study species that exhibit little or no female song, studies on more taxa with natural female song are needed to evaluate the natural association of female song and testosterone (Rose et al., 2022).

In this study, we focus on the northern house wren (*Troglodytes aedon*), a temperate-breeding species with naturally occurring female song (Johnson and Kermott, 1990; Krieg and Getty, 2016). Northern house wrens are common temperate-breeding songbirds with sexually dimorphic songs (Keck et al., *in press;* Krieg and Wade, 2023). Compared with males, female house wrens have very low plasma testosterone and smaller volumes of HVC, Area X, and RA than males but exhibit seasonal neurogenesis in HVC (Krieg and Wade, 2023; Odom, unpublished data). Female house wrens sing reliably in the early breeding season through egg-laying with female song steeply declining at incubation (Krieg and Getty, 2016; Odom, unpublished data). Female songs, on average, are less complex than male songs, and individuals vary wildly in song structure and output, with some birds singing very similar songs to males (Johnson and Kermott, 1990; Krieg and Getty, 2016; Krieg and Wade, 2023). In female house wrens, volumes of RA and Area X correlate with song complexity, suggesting that song control regions regulate song structure in similar ways to males (Alward et al., 2017; Ball and Balthazart, 2020; Krieg and Wade, 2023).

As a first step to investigate testosterone sensitivity in female house wren song nuclei, we quantified the expression of androgen receptor (AR), estrogen receptors (ERα and ERβ), and aromatase (AROM) in HVC, Area X, and RA. In the brain, testosterone and its metabolite 5α-dihydrotestosterone can bind to AR, inducing its effects, or aromatase can convert testosterone to 17β-estradiol, which can bind either of two estrogen receptors, ERα and ERβ. AR, ERα, and ERβ can then activate transcription factors that regulate gene expression (Schlinger, 1997). In the song control system, AR is expressed in HVC and RA, and in some species Area X; ERα is expressed in HVC and sometimes around the dorsal aspect of RA (Gahr 2014; Rose et al. 2022); ERβ and aromatase are generally not expressed in song nuclei in the species and regions that have been examined but are less well studied (Bernard et al., 1999; Frankl-Vilches and Gahr, 2018; Metzdorf et al., 1999). In other species, neuroanatomical distribution of AR and ERα is similar in males and females, although males have higher AR expression especially in HVC in certain species, particularly zebra finch (Gahr, 2014; Rose et al. 2022). Changes in AR and ERα expression can modulate hormone sensitivity (Gahr and Metzdorf, 1997), so we hypothesized that upregulation of steroid receptors in females could compensate for low testosterone. In this study, we quantified mRNA levels of the steroid receptors AR, ERα, ERβ, and AROM in the song nuclei HVC, Area X, and RA of the house wren. We then evaluated whether expression of these transcripts varies between sex, season, and brain region in this temperate-breeding songbird with female song.

## METHODS

### Field collection

We performed this study on adult house wrens across four different breeding stages: nonbreeding, pre-laying, egg laying, and incubation. In male house wrens, song and testosterone are low in the nonbreeding season; both increase dramatically in the early breeding season and then steadily decrease throughout the breeding cycle (Cramer, 2012; Johnson, 2020; Odom et al., *in prep*). Similarly, female house wren song peaks during the pre-laying to egg-laying period, then rapidly decreases at incubation, although female house wren circulating testosterone is low and does not change seasonally with female song (Krieg and Getty, 2016; Odom et al. in prep). Nonbreeding male house wrens were caught in February 2022 in Alachua County, Florida. Breeding male and female house wrens were caught from May 2022 to July 2022 and May 2023 to July 2023 in Monroe County and Wayne County, Pennsylvania. We collected five to six wrens of each sex per breeding stage for a total of n = 22 males and n = 18 females. Nonbreeding females were not included in this study because we did not find any at our nonbreeding study site.

All house wrens were caught in mist nets using playback of male or female song as a lure. All individuals were then rapidly decapitated, and the whole brain was quickly removed. Each brain was bisected on ice using a clean razor blade. The left hemisphere of each wren was immediately flash frozen on powdered dry ice and stored at −80°C for RNA extraction and sequencing. The right hemisphere was formaldehyde-fixed for 24 hours, stored in 30% sucrose in PBS for 1 to 3 days, frozen on dry ice, then stored at −80°C for histological preparation. In the field, most brains were processed (flash frozen or immersed in fixative) within 7 to 11 minutes after decapitation, except for a few individuals across treatments that took up to 15 minutes to process.

### Tissue sample preparation

The flash frozen left hemispheres of house wren brains were stored at −80°C for between 3 to 18 months before preparation for RNA extraction. Within a month prior to RNA extraction, all brains were sectioned and punched (microdissected) according to Palkovit’s punch technique (e.g., George and Rosvall, 2025; Heimovics et al., 2012; Palkovits, 1983).

To collect punches, brains were first sectioned in the coronal plane at 200 μm on a cryostat at −16 to −18°C. Sections were thaw-mounted on microscope slides, and all microdissections were made while in the cryostat at −25 to −20°C. Punches of Area X and RA were made using a 1.0 mm (inner diameter) stainless-steel sample corer, whereas HVC was punched as three lateral to medial punches using a 0.5 mm (inner diameter) stainless-steel sample corer (catalogue #18035-50; Fine Science Tools, Foster City, CA, USA). All punches were stored in 1.5 mL microcentrifuge tubes at −80°C until RNA extraction.

The specific sections and locations to punch were determined using Nissl and Hu-stained tissue from the formaldehyde-fixed hemisphere of each brain. In these stained tissues, we identified the song control regions of interest and calculated the average distance (number of sections) from major landmarks including TrSM split, anterior commissure (CoA), and CoA + Tractus occipito-mesencephalicus (OM). Specifically, we punched each region over four 200 μm sections, spanning the majority of each of the respective song nucleus. Each nucleus was punched within the following sections based on the following landmarks: Area X was punched caudal to rostral starting three to four sections rostral to TrSM split, HVC was punched rostral to caudal starting two sections caudal to CoA + OM, and RA was punched rostral to caudal starting six sections caudal to CoA + OM (which is equivalent to the end of HVC in house wrens). Though we carefully determined the precise locations to punch, female song nuclei are very small, so the punched area may have been larger in area than the entire song nucleus (Krieg and Wade, 2023). It is thus possible that our tissue punches contained non-target tissue, especially for smaller nuclei such as HVC and Area X in females and RA in both sexes.

### RNA extraction & RT-qPCR

We extracted total RNA from tissue punches using TRizol reagent according to the manufacturer’s protocol (Invitrogen, Carlsbad, CA). This RNA was treated for leftover genomic DNA with DNAse I (New England Biolabs, Ipswich, MA). RNA was then purified using the RNeasy MinElute Cleanup Kit (Qiagen, Hilden, DE) according to the manufacturer’s protocol. Concentration and purity of the resulting RNA extract was measured with a NanoDrop One Spectrophotometer (Thermo Fischer Scientific, Waltham, MA; Table S1). The RNA was stored at −80°C until later use.

We reverse transcribed total RNA into cDNA using the SuperScript IV First-Strand Synthesis System (Thermo Fisher Scientific) according to the manufacturer’s protocol. Volumes of each sample were adjusted according to their RNA concentration to add the same mass of RNA to each reaction. We separately prepared negative RT controls for each sample, in which reverse transcriptase was replaced with an equal volume of water.

We adapted gene specific primers for AR, AROM, ERα, and ERβ. Primers for AR, AROM, and ERα were adapted from previous studies to better match the melting temperature for our protocol (see Table 1). Primers for ERβ were designed to span the final intron of the Zebra Finch (*Taeniopygia guttata)* ERβ gene (NCBI Reference Sequence: XM_030275146.3), producing a 101 bp amplicon. We verified the sequences complemented across passerines. Primers for AR, AROM, ERα, and ERβ were obtained commercially (Integrated DNA Technologies, Coralville, IA).

**Table 1.**
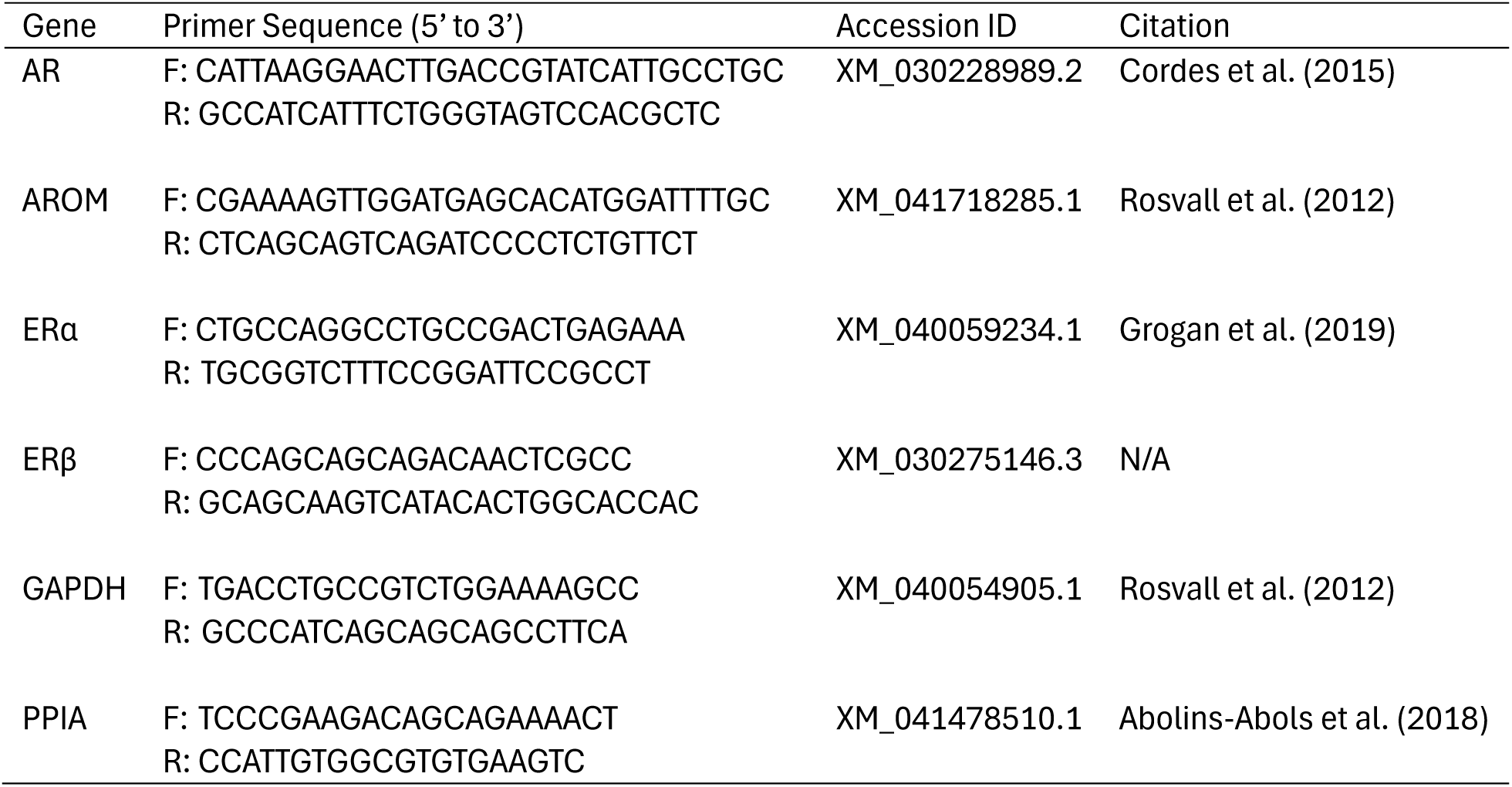
Primer sequences. AR: androgen receptor, AROM: aromatase, ERα: estrogen receptor α, ERβ: estrogen receptor β, GAPDH: glyceraldehyde-3-phosphate dehydrogenase, PPIA: peptidylprolyl isomerase A.

We selected GAPDH and PPIA as reference genes because this combination of genes was identified as stable in the brain of two other songbird species (Zinzow-Kramer et al., 2015). We confirmed these genes were stably expressed in our samples using geNorm (Vandesompele et al., 2002) and that expression did not vary across sex, brain region, or breeding stage using NormFinder (Andersen et al., 2004). Primers for reference genes were obtained from the references in Table 1.

We performed qPCR on a CFX96 Touch Real-Time PCR Detection System (Bio-Rad, Hercules, CA). All genes for an individual were run on the same plate, and samples were run in a balanced design with an equal number of male and female samples on each plate. Each sample was run in triplicate alongside a negative RT control. A single reaction of 10 μL consisted of 1 μL iTaq Universal SYBR Green Supermix (Bio-Rad), 1 μL each of the appropriate forward and reverse primer (10 μM), 1 μL cDNA, and 6 μL RNase-free water. Because our experiment required multiple plates, we also added an inter-plate calibrator to each plate, which came from pooled cDNA from Area X of three nonbreeding males not included in the study. The thermocycler began with a starting denaturation cycle for 30 seconds at 95°C, followed by 40 cycles of 15 seconds at 95°C, 30 seconds at 59°C, then a plate reading. The program then performed a melt curve analysis with 0.5°C increments every 2 to 5 seconds from 65°C-95°C.

We quantified the expression of AR, AROM, ERα, and ERβ relative to the reference genes GAPDH and PPIA. Using the CFX Maestro Software (Bio-Rad), we adjusted for differences in running conditions of individual plates using the inter-plate calibrator. The cycle threshold (Ct) values of technical replicates were averaged. We then averaged Ct values of the reference genes GAPDH and PPIA and used this value to calculate ΔCt (Ct [gene of interest] – Ct [reference genes]). ΔCt values were plotted and used for all analyses.

### Statistical analyses

We performed all statistical analyses in R v 4.4.1 (R Core Team, 2019). To analyze the effects of brain region, sex, and breeding stage on gene expression, we built linear mixed models (LMMs) with quantification data for each gene using the “lme4” package in R. Models included brain region, sex, and breeding stage as fixed effects and bird identity as a random effect. In addition, we analyzed the effects of sex and breeding stage within each brain region using linear models (LMs) for each combination of gene and brain region. These models included sex and breeding stage as fixed effects with no random effects. We initially tested interaction terms but later removed them because they were non-significant.

We assessed normality and heteroscedasticity of each model to ensure the distribution of the data did not violate model assumptions. Only the LM for ERα in HVC departed from normality of residuals, so we performed a reciprocal transformation on its data.

Significance of fixed effects for both LMMs and LMs were analyzed using type II ANOVAs; for LMMs specifically, we used the “lmerTest” package. We performed post hoc Tukey’s tests to assess which variables contributed to significant differences using the “emmeans” package to estimate marginal means. We accepted an alpha-level of p = 0.05 as statistically significant for all analyses.

### Ethics Statement

All work was conducted in accordance with the Association for the Study of Animal Behaviour’s guidelines and the Ornithological Council’s Guidelines to the Use of Wild Birds in Research. Our methods were approved by the Institutional Animal Care and Use Committee (IACUC; #FR-APR-21-17), University of Maryland, College Park. These activities were permitted under Federal U.S. Fish and Wildlife Service Scientific Collecting Permit number MB01550B, Pennsylvania State Game and Conservation Commission Scientific Collecting Permit number 52256, Pennsylvania State Game and Conservation Commission Specific Use Permit number 52524, and Florida Fish and Wildlife Conservation Commission Scientific Collecting Permit number LSSC-12-00016C.

## RESULTS

### Brain Region Differences

Brain region had a significant effect on all genes (p < 0.001 to p = 0.006, Table 2, Figure 1). Post hoc Tukey’s tests revealed that HVC, RA, and Area X all significantly differed from each other, again for all genes (p < 0.001, Table S2). AR, AROM, and ERα receptor expression was highest in HVC, then RA, then Area X. In ERβ, expression was highest in RA, then HVC, then Area X (Table 2, Figure 1).

**Figure 1.**
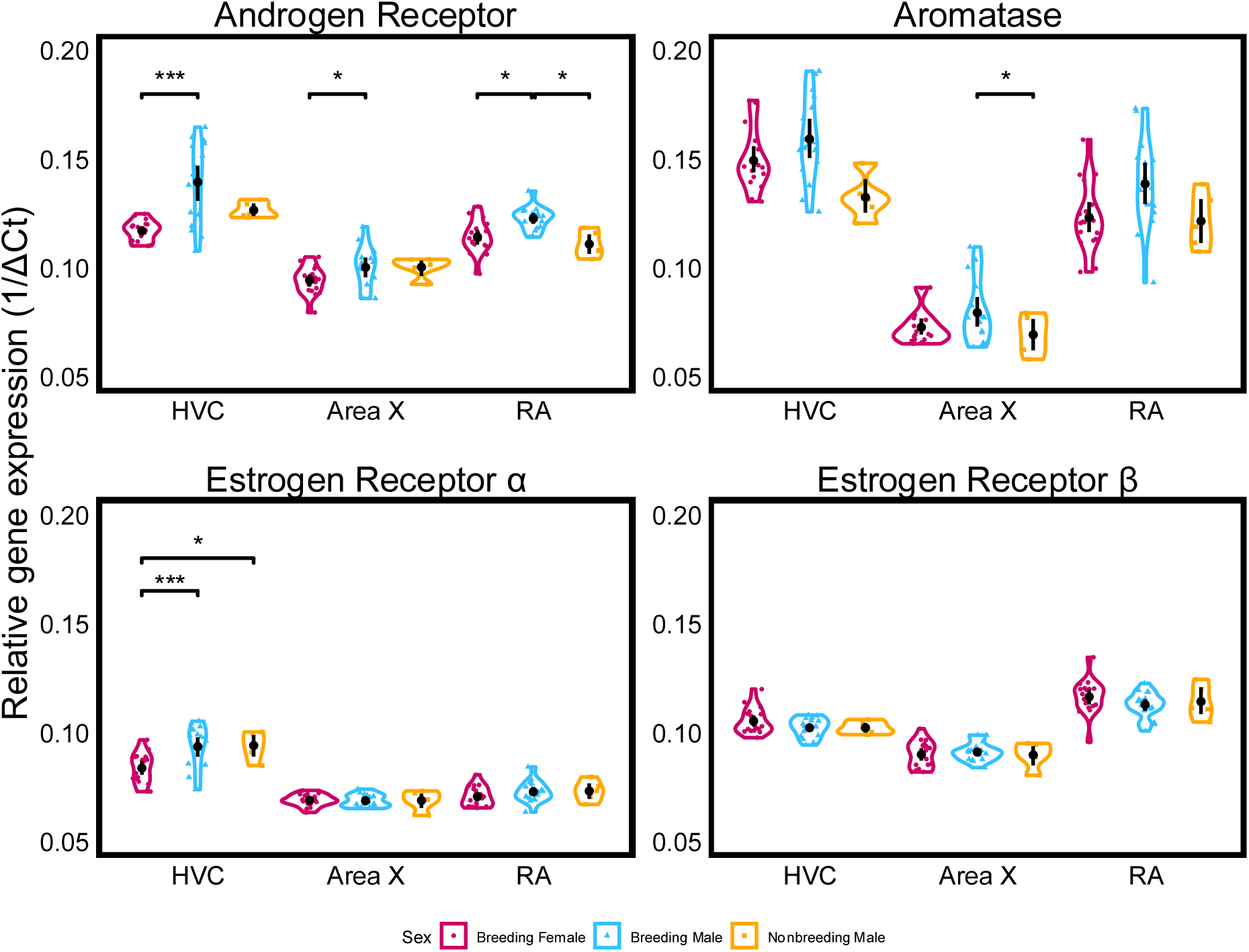
mRNA of steroid receptors is not expressed at greater levels in breeding female northern house wrens versus breeding males. Gene expression of target genes is relative to the average Ct of the housekeeping genes GAPDH and PPIA. All brain regions are significantly different from each other. Only significant comparisons between sexes are shown. HVC: acronym serves as proper name, RA: robust nucleus of the arcopallium.

**Table 2.** Summaries of linear mixed models evaluating effects of brain region along with breeding stage and sex on steroid receptor gene expression. HVC: acronym serves as proper name, RA: robust nucleus of the arcopallium.

| Gene | Predictor variable | Estimate | Std. Error | df | p-value |  |
| --- | --- | --- | --- | --- | --- | --- |
| AR | **Sex: Male** | -0.82 | 0.14 | 35 | **<0.001** | *** |
|  | Breeding Stage: Nonbreeding | 0.35 | 0.24 | 35 | 0.153 |  |
|  | Breeding Stage: Egg laying | -0.23 | 0.18 | 35 | 0.211 |  |
|  | Breeding Stage: Incubating | -0.11 | 0.18 | 35 | 0.522 |  |
|  | **Brain Region: HVC** | -2.37 | 0.15 | 78 | **<0.001** | *** |
|  | **Brain Region: RA** | -1.75 | 0.15 | 78 | **<0.001** | *** |
| AROM | **Sex: Male** | -0.69 | 0.30 | 35 | **0.026** | * |
|  | **Breeding Stage: Nonbreeding** | 1.16 | 0.50 | 35 | **0.025** | * |
|  | Breeding Stage: Egg laying | -0.24 | 0.36 | 35 | 0.521 |  |
|  | Breeding Stage: Incubating | -0.12 | 0.36 | 35 | 0.742 |  |
|  | **Brain Region: HVC** | -6.93 | 0.25 | 78 | **<0.001** | *** |
|  | **Brain Region: RA** | -5.70 | 0.25 | 78 | **<0.001** | *** |
| ERα | **Sex: Male** | -0.53 | 0.18 | 35 | **0.007** | ** |
|  | Breeding Stage: Nonbreeding | -0.17 | 0.31 | 34 | 0.592 |  |
|  | Breeding Stage: Egg laying | -0.01 | 0.23 | 35 | 0.964 |  |
|  | Breeding Stage: Incubating | -0.26 | 0.23 | 34 | 0.251 |  |
|  | **Brain Region: HVC** | -3.23 | 0.21 | 77 | **<0.001** | *** |
|  | **Brain Region: RA** | -0.59 | 0.21 | 77 | **0.006** | ** |
| ERβ | Sex: Male | 0.14 | 0.11 | 35 | 0.213 | |
|  | Breeding Stage: Nonbreeding | -0.16 | 0.18 | 35 | 0.381 |  |
|  | Breeding Stage: Egg laying | -0.16 | 0.13 | 35 | 0.239 |  |
|  | **Breeding Stage: Incubating** | -0.36 | 0.13 | 35 | **0.011** | * |
|  | **Brain Region: HVC** | -1.44 | 0.12 | 78 | **<0.001** | *** |
|  | **Brain Region: RA** | -2.36 | 0.12 | 78 | **<0.001** | *** |

p-values calculated through package lmerTest.
Bolded values are statistically significant based on a $p$ -value of 0.05.
Significance codes: '\*\*\*' $0 \leq p \leq 0.001$ | '\*\*' $0.001 < p \leq 0.01$ | '\*' $0.01 < p \leq 0.05$ | '.' $0.05 < p \leq 0.1$

### Sex Differences

Sex had a significant effect in AR, AROM, and ERα but not ERβ (p < 0.001 to p = 0.02, Table 2, Figures 1 & 2). When sex differences occurred, females always had lower expression than males. Table 3 shows which brain regions had sex differences: for AR, expression was higher in males than females in all three brain regions; for AROM, expression was higher in males only in Area X; for ERα, expression was higher in males only in HVC; ERβ did not differ between sex in any region.

**Figure 2.**
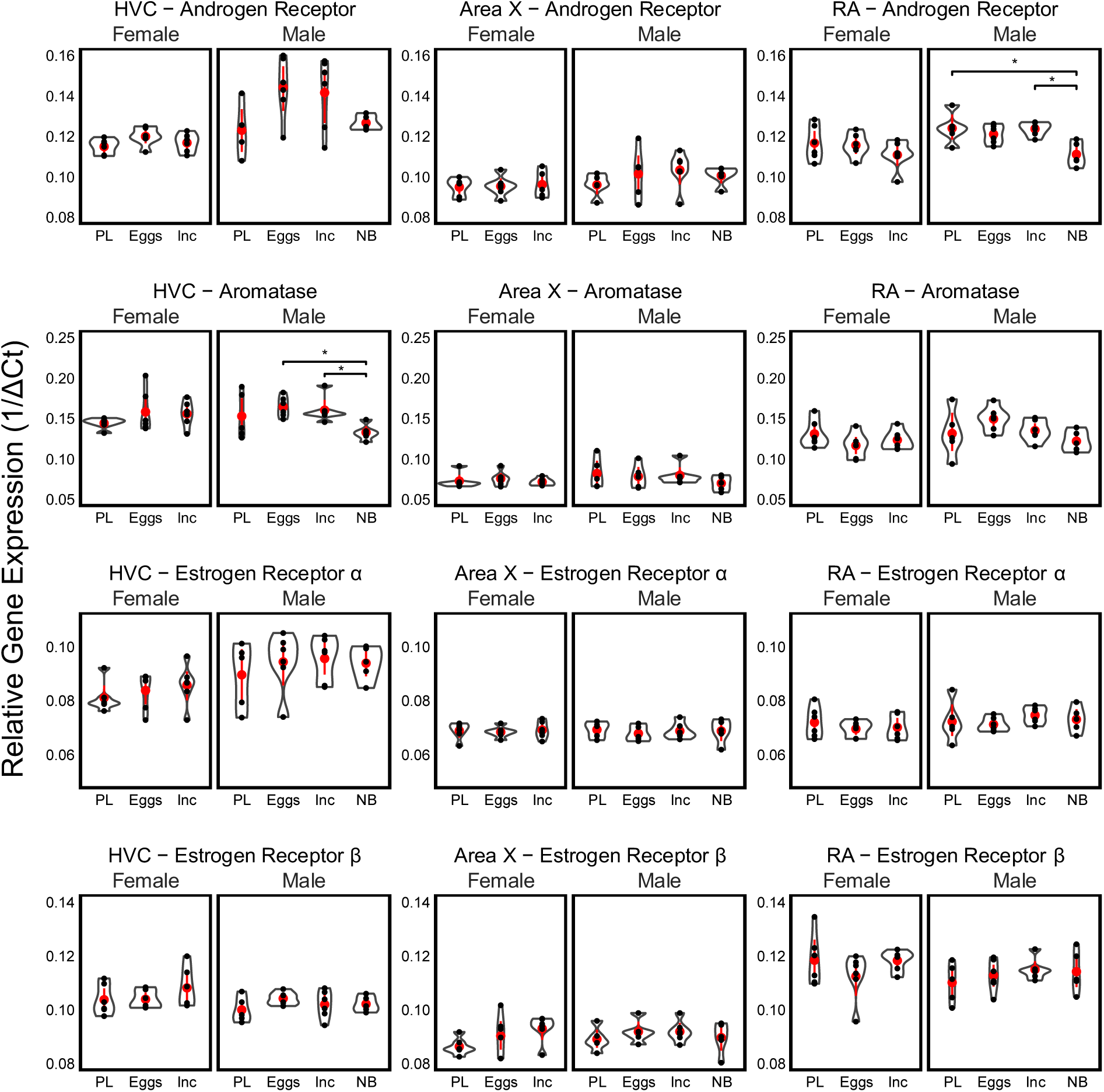
Relative mRNA expression of steroid receptors for each combination of gene, brain region, sex, and breeding stage. Gene expression of target genes is relative to the average Ct of the housekeeping genes GAPDH and PPIA. PL: pre-laying, Eggs: egg laying, Inc: incubation, NB: nonbreeding. Only significant comparisons are shown.

**Table 3.**
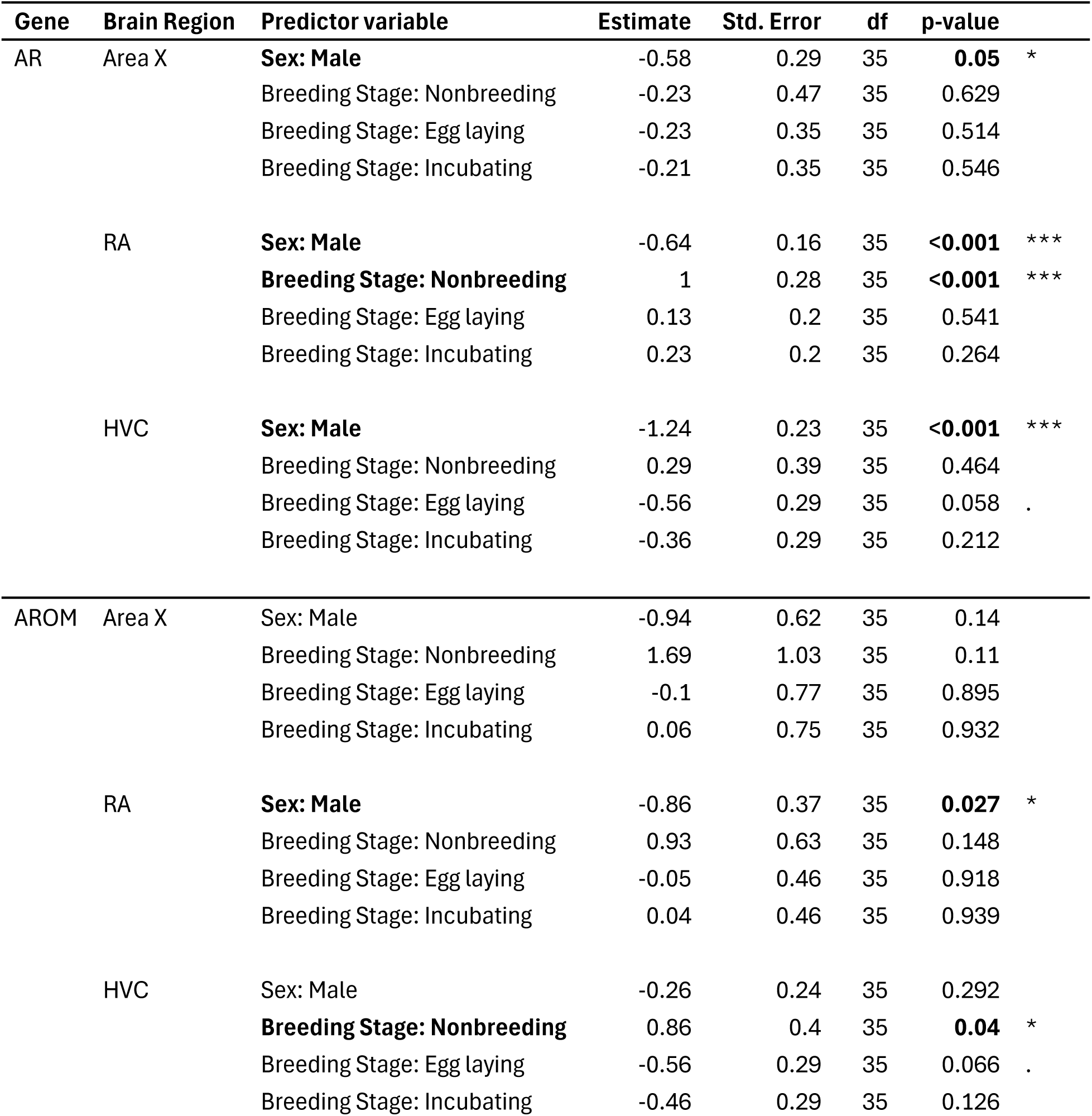
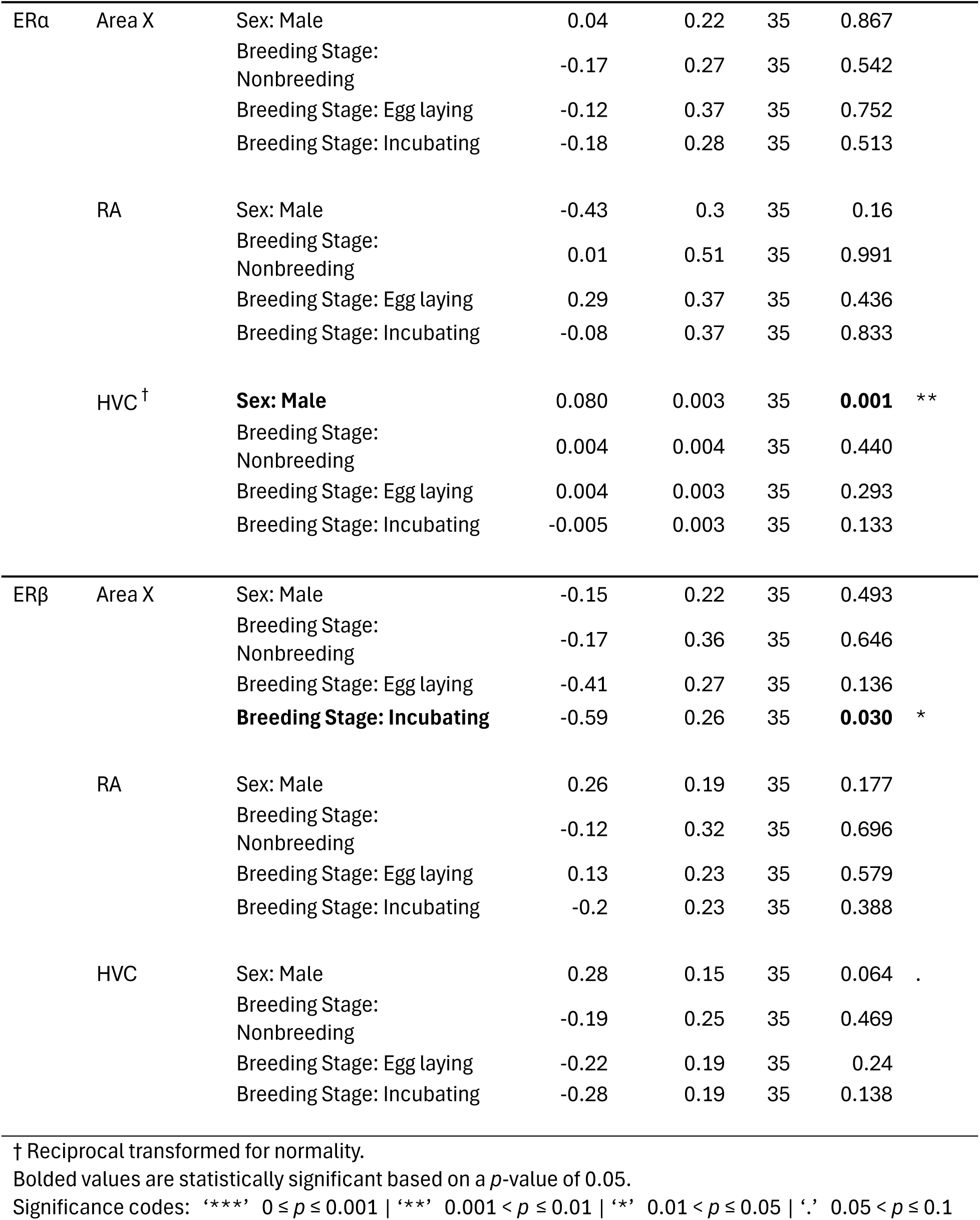
Summaries of linear models evaluating effects of breeding stage and sex on steroid receptor gene expression within brain regions.

| Gene | Brain Region | Predictor variable | Estimate | Std. Error | df | p-value |  |
| --- | --- | --- | --- | --- | --- | --- | --- |
| AR | Area X | **Sex: Male** | -0.58 | 0.29 | 35 | **0.05** | * |
|  |  | Breeding Stage: Nonbreeding | -0.23 | 0.47 | 35 | 0.629 |  |
|  |  | Breeding Stage: Egg laying | -0.23 | 0.35 | 35 | 0.514 |  |
|  |  | Breeding Stage: Incubating | -0.21 | 0.35 | 35 | 0.546 |  |
|  | RA | **Sex: Male** | -0.64 | 0.16 | 35 | **<0.001** | *** |
|  |  | **Breeding Stage: Nonbreeding** | 1 | 0.28 | 35 | **<0.001** | *** |
|  |  | Breeding Stage: Egg laying | 0.13 | 0.2 | 35 | 0.541 |  |
|  |  | Breeding Stage: Incubating | 0.23 | 0.2 | 35 | 0.264 |  |
|  | HVC | **Sex: Male** | -1.24 | 0.23 | 35 | **<0.001** | *** |
|  |  | Breeding Stage: Nonbreeding | 0.29 | 0.39 | 35 | 0.464 |  |
|  |  | Breeding Stage: Egg laying | -0.56 | 0.29 | 35 | 0.058 | . |
|  |  | Breeding Stage: Incubating | -0.36 | 0.29 | 35 | 0.212 |  |
| AROM | Area X | Sex: Male | -0.94 | 0.62 | 35 | 0.14 |  |
|  |  | Breeding Stage: Nonbreeding | 1.69 | 1.03 | 35 | 0.11 |  |
|  |  | Breeding Stage: Egg laying | -0.1 | 0.77 | 35 | 0.895 |  |
|  |  | Breeding Stage: Incubating | 0.06 | 0.75 | 35 | 0.932 |  |
|  | RA | **Sex: Male** | -0.86 | 0.37 | 35 | **0.027** | * |
|  |  | Breeding Stage: Nonbreeding | 0.93 | 0.63 | 35 | 0.148 |  |
|  |  | Breeding Stage: Egg laying | -0.05 | 0.46 | 35 | 0.918 |  |
|  |  | Breeding Stage: Incubating | 0.04 | 0.46 | 35 | 0.939 |  |
|  | HVC | Sex: Male | -0.26 | 0.24 | 35 | 0.292 |  |
|  |  | **Breeding Stage: Nonbreeding** | 0.86 | 0.4 | 35 | **0.04** | * |
|  |  | Breeding Stage: Egg laying | -0.56 | 0.29 | 35 | 0.066 | . |
|  |  | Breeding Stage: Incubating | -0.46 | 0.29 | 35 | 0.126 |  |
| ERα | Area X | Sex: Male | 0.04 | 0.22 | 35 | 0.867 | |
|  |  | Breeding Stage: Nonbreeding | -0.17 | 0.27 | 35 | 0.542 |  |
|  |  | Breeding Stage: Egg laying | -0.12 | 0.37 | 35 | 0.752 |  |
|  |  | Breeding Stage: Incubating | -0.18 | 0.28 | 35 | 0.513 |  |
|  | RA | Sex: Male | -0.43 | 0.3 | 35 | 0.16 |  |
|  |  | Breeding Stage: Nonbreeding | 0.01 | 0.51 | 35 | 0.991 |  |
|  |  | Breeding Stage: Egg laying | 0.29 | 0.37 | 35 | 0.436 |  |
|  |  | Breeding Stage: Incubating | -0.08 | 0.37 | 35 | 0.833 |  |
|  | HVC<sup>†</sup> | **Sex: Male** | 0.080 | 0.003 | 35 | **0.001** | ** |
|  |  | Breeding Stage: Nonbreeding | 0.004 | 0.004 | 35 | 0.440 |  |
|  |  | Breeding Stage: Egg laying | 0.004 | 0.003 | 35 | 0.293 |  |
|  |  | Breeding Stage: Incubating | -0.005 | 0.003 | 35 | 0.133 |  |
| ERβ | Area X | Sex: Male | -0.15 | 0.22 | 35 | 0.493 | |
|  |  | Breeding Stage: Nonbreeding | -0.17 | 0.36 | 35 | 0.646 |  |
|  |  | Breeding Stage: Egg laying | -0.41 | 0.27 | 35 | 0.136 |  |
|  |  | **Breeding Stage: Incubating** | -0.59 | 0.26 | 35 | **0.030** | * |
|  | RA | Sex: Male | 0.26 | 0.19 | 35 | 0.177 |  |
|  |  | Breeding Stage: Nonbreeding | -0.12 | 0.32 | 35 | 0.696 |  |
|  |  | Breeding Stage: Egg laying | 0.13 | 0.23 | 35 | 0.579 |  |
|  |  | Breeding Stage: Incubating | -0.2 | 0.23 | 35 | 0.388 |  |
|  | HVC | Sex: Male | 0.28 | 0.15 | 35 | 0.064 | . |
|  |  | Breeding Stage: Nonbreeding | -0.19 | 0.25 | 35 | 0.469 |  |
|  |  | Breeding Stage: Egg laying | -0.22 | 0.19 | 35 | 0.24 |  |
|  |  | Breeding Stage: Incubating | -0.28 | 0.19 | 35 | 0.138 |  |

† Reciprocal transformed for normality.
Bolded values are statistically significant based on a $p$ -value of 0.05.
Significance codes: '\*\*\*' $0 \leq p \leq 0.001$ | '\*\*' $0.001 < p \leq 0.01$ | '\*' $0.01 < p \leq 0.05$ | '.' $0.05 < p \leq 0.1$

### Breeding Stage Differences

We found breeding stage differences in AR, AROM, and ERα. Our analyses were complicated because we did not have any nonbreeding females in our study, so we were unable to compare nonbreeding females with other groups. Instead, we compared breeding females, breeding males, and nonbreeding males (Figures 1 & 2, Table S3). Breeding stage differences were often from breeding females having significantly lower expression than breeding males, which was especially pronounced for AR expression in HVC (p < 0.001, Table S3, Figures 1 & 2). Interestingly, breeding female ERα expression was significantly lower than even nonbreeding males in HVC (p = 0.01).

Nonbreeding males had significantly lower expression than breeding males in AROM in HVC (p = 0.005) and AR in RA (p = 0.008, Figure 2). This suggests that males may increase expression of these genes from the nonbreeding season to the breeding season. However, we did not find that expression differed between the three stages within the breeding season. In all other genes and brain regions, expression was not significantly different between the nonbreeding, pre-laying, egg laying and incubation stages (Table S3).

## DISCUSSION

We found that AR expression in female house wrens was lower than in breeding males—and even some nonbreeding males—in all three brain regions. This does not support for our initial hypothesis that females might have increased neural AR in song control regions as a mechanism to increase sensitivity to low levels of circulating testosterone. In addition, we did not see an elevation of female AR expression in early breeding stages when females sing most, suggesting AR expression does not shift seasonally along with female song production. Furthermore, we saw no evidence that females had increased sensitivity to circulating aromatized androgens: ERα was lower in females than males, and ERβ expression was similar between the sexes.

We also found interesting seasonal and individual differences in expression, especially in males. Most pronounced was an increase in male AR expression in the breeding season compared with the nonbreeding season, especially in RA (although a similar, though non-significant trend was found in HVC). AROM expression was also greater in the breeding season compared to the nonbreeding season in Area X. AR and AROM expression varied greatly among individual breeding males, specifically in HVC, to the extent that some breeding males had lower expression than nonbreeding males. Therefore, while singing female house wrens do not appear to compensate for low circulating testosterone by increasing neural steroid sensitivity beyond levels in males, male house wrens do increase their AR expression and likely their sensitivity during the breeding season. Our findings in breeding males is congruent with previous findings that testosterone, AR and other receptor expression, and even aromatase play an important role in mediating seasonal and individual differences in male singing behavior (Alward et al., 2016, 2017; Rose et al., 2022). Few studies have examined sex differences in ER and AROM (Rose et al., 2022), so we provide a valuable contribution to what is known about these molecules, especially in this non-model, wild-breeding species.

### Sex Differences in Steroid Sensitivity: Implications on Female Song

Prior studies demonstrate different degrees of sex differences in AR expression, from females having less to similar quantities of AR compared with males (Gahr 2014; Rose et al., 2022). However, no previous studies report females having greater AR expression in song control nuclei than males (Rose et al., 2022). Here, we also find that naturally singing female house wrens have lower AR expression than males. Previous studies have found that female zebra finch have fewer AR expressing neurons (Arnold and Saltiel, 1979; Gahr, 2014; Gahr and Metzdorf, 1997; Rose et al., 2022). In addition, the larger volumes of HVC, RA, and Area X likely compound this sex difference: song control region volumes in male house wrens are two to four times larger than in females (Krieg and Wade, 2023). Thus, female house wrens likely have fewer total neurons in HVC and fewer neurons expressing AR.

This male-bias in steroid receptors and steroid processing enzymes matches the male-bias in singing behavior. Compared with males, female house wrens sing much less frequently and, on average, have shorter, simpler, and less stereotyped songs (Johnson and Kermott, 1990; Krieg and Getty, 2016; Krieg and Wade, 2023). Female house wren songs also vary considerably between individuals. In male songbirds, testosterone action in HVC and RA regulates song complexity and stereotypy (Alward et al., 2017); therefore, in female house wrens, low testosterone concentrations in concert with similar or reduced testosterone sensitivity in HVC and RA may result in simple, variable songs.

However, female song peaks during specific times (e.g. beginning of the egg-laying stage) and specific social contexts (e.g. conspecific conflicts, redirecting mate attention; Johnson and Kermott, 1990; Krieg and Getty, 2016). Steroid receptor expression in other parts of the brain can change rapidly in response to social context (Fuxjager et al., 2010). By grouping female samples into broad breeding stages, it is possible we missed subtle changes in steroid receptor expression that support female song production. Even if this was the case, if female song is largely regulated by steroid receptor density, we would have expected to see reduced expression during incubation when females sing the least, which we did not.

### Alternate Mechanisms if Female Song is Less Dependent on Plasma Testosterone

If female song is less dependent on circulating plasma testosterone, what other mechanisms might regulate female singing? One possibility is that testosterone could be synthesized in the brain separately from gonadal testosterone (e.g., localized production of neurosteroids; Remage-Healey et al., 2010). In male songbirds, when circulating testosterone is low during the nonbreeding season, local conversion of DHEA to testosterone in the brain occurs to regulate aggressive behavior (Soma et al., 2015). If regulated by similar mechanisms, female song might still be steroid-mediated even though circulating plasma testosterone is low. Nevertheless, our results, combined with other studies, indicate that female songbirds have lower AR expression than breeding males in these major song control regions; therefore, song control regions of female house wrens and other female songbirds are likely less sensitive than males to testosterone; thus, this mechanism is unlikely to be the sole regulator of female song.

Alternatively, testosterone may be acting outside of the primary song control nuclei to modulate female song. In the canary’s medial preoptic nucleus (POM), a region embedded in the preoptic area, testosterone plays a large role in mediating the motivation to sing, although not song quality (Alward et al., 2013). As such, while testosterone in HVC and RA specifically may control aspects such as song stereotypy or performance in males (Alward et al., 2014; Alward et al., 2017), testosterone’s effect of increasing motivation to sing, relevant in males and naturally singing females, may be found within the POM. Few studies have investigated sex differences in POM in songbirds, but in Japanese quail, a non-songbird, POM volume is greater in males (Ball and Balthazart, 2020; Panzica et al., 1996).

A third possibility is that hormones, primarily estradiol, and molecular mechanisms other than testosterone could more directly regulate female bird song. Aromatase in the brain acts locally to convert testosterone to estradiol in males via aromatase action (Schlinger, 1997; Vandries et al., 2019). In this way, both male and female song might be primarily regulated by estradiol. This is an intriguing option because estrogenic signaling by aromatized testosterone appears to be a main mechanism regulating temporary increases in song in male canaries (Alward et al., 2016). In addition, Heimovics et al. (2012) show estradiol signaling in the brain regulates seasonal changes in aggressive behavior in male song sparrows. Moreover, song production and song complexity can be increased in female canaries by administering estrogens (Vandries et al., 2019). However, in both males and females, estrogen receptor expression is low and not nearly as widely distributed as AR in the song control system of songbirds (Gahr, 2014; Rose et al., 2022).

A final possibility is that female song in house wrens is governed mechanisms and changes in gene expression that are not testosterone or steroid-mediated. While this seems unlikely, Goymann and Wingfield (2014) argue that the evidence for testosterone mediating female song is still lacking. Since female song is still not well understood in wild songbird populations with natural female song, more work is needed to investigate the role of testosterone, estradiol, and the downstream molecular pathways involved in natural birdsong in a range of species with variation in female song.

### Temporal Changes in Expression of Steroid Receptors

While female steroid receptor expression did not change significantly across season, we saw interesting seasonal changes in male steroid receptor expression. We saw increased expression in AR in RA and AROM in HVC during the breeding season compared with the nonbreeding season. AR and AROM expression also increased in the breeding season and decreased during the nonbreeding season in male canaries (Fusani et al., 2000). Together, these results are consistent with testosterone-mediated regulation of male song, with heightened sensitivity in the breeding season.

Also of note were the large individual variations in expression, especially of AR in HVC in breeding males. This is especially interesting because testosterone action in the HVC of songbirds specifically regulates variation in syllable usage and sequencing (i.e., “complexity”; Alward et al. 2017). Therefore, variation in AR receptor expression in HVC could directly impact or regulate individual variation in male song quality during the breeding season. As breeding male birdsong complexity is largely thought of as a quality indicator that impacts female mating decisions, the high individual variation of AR expression in HVC could translate to phenotypic variation in song that leads to variation in mating success. In addition, such individual differences in testosterone sensitivity may be important for individual variation in sexual behavior, aggression, and competition. Individual variation in aggression has been linked to individual variation in AR expression and sensitivity in other songbird species (Rosvall et al., 2012). Further research is needed to determine if this is the case in house wrens.

### Expression of Estrogen Receptors and Aromatase

ERα was expressed at relatively low levels in HVC, and at very slight levels in Area X and RA. This matches results in other species which report ERα in HVC and dorsal RA using in-situ hybridization (Gahr, 2014). Estrogen in the song control system has been implicated in song learning, and supplementary estrogen can increase the volume of the song nuclei similarly to testosterone (Schlinger, 1997; Tramontin et al., 2003). Estrogen may also have specific effects such as increasing motivation to sing, temporarily or long-term (Alward et al., 2016).

We detected relatively high AROM and ERβ expression in both sexes in most song control regions, contrary to studies in other songbirds. Previous studies have found little to no aromatase in song control regions (Gahr, 2014). Previous studies have also not found ERβ expression in the song control system (Bernard et al., 1999; Rose et al. 2022), except Ko et al. (2021), which found ERβ in HVC of female canaries. However, studies investigating the distribution of aromatase or estrogen receptors in female song control regions are few. Our detections of AROM and ERβ may differ from previous studies due to our use of qPCR. qPCR is more sensitive than in-situ hybridization, the more common approach of prior research, since it involves multi-fold amplification of the genes of interest. Therefore, we may have amplified very low densities of AROM and ERβ found in locations it is previously undetected; for example, within synaptic terminals of the song system (Remage-Healey et al. 2010). Alternatively, our tissue punches could have included adjacent non-target tissue that contained ERβ and AROM. Specifically, the NCM directly ventral to HVC and dorsal-rostral to RA expresses both these genes (Gahr et al., 2014; Rose et al., 2022), A disadvantage of our approach is that we were not able to localize receptor expression. Therefore, we recommend follow up studies using localization methods to determine the exact distribution of these genes in male and female house wren song control regions.

## Conclusion

Taken together, our study suggests that female house wrens do not compensate for lower testosterone concentrations by increased neural sensitivity to steroid hormones alone: none of AR, AROM, ERα, or ERβ were expressed in greater levels in song control regions of female house wrens than males. We found expression of AR and AROM increases from the nonbreeding season to the breeding season in males, but we did not find significant differences in expression in females within the breeding season. Our study provides a unique example of a wild species with natural song in both sexes, the house wren; we recommend future studies also include a wider selection of species in which females sing to address the many gaps in understanding of female song in birdsong research.

## Acknowledgements

We are grateful to Ailin Dolson-Fozio, Maxine Hsu, and Sarah Keck for help with breeding house wren fieldwork in Pennsylvania, and we thank the Pennsylvania State Game Commission for assistance with permits and logistics of land use. We thank the following faculty, staff, and students at the University of Florida for assistance with nonbreeding field research: Scott Robinson, Andrew Kratter, Liz Hurtado, Grant Terrell, and Rachael Woods. We thank Kim Rosvall for guidance on primers, reference genes, and feedback on project design. We also thank members of the Ball-Dooling lab for feedback throughout project planning and analysis. We acknowledge the original inhabitants and stewards of the Land: the Lenape, the Munsee, the Shawnee, and the Susquehannock. We thank the University of Maryland, College Park and the University of Scranton for funding and logistical support. We thank the University of the Pacific for internal funding in the form of a Pre-Tenure Award to Karan Odom and a Fred and Marguerite Early Award to John Yoo.

## Notes

### Competing Interest Statement

The authors have declared no competing interest.

