## Supplementary Material for "Breeding males, but not females, have elevated androgen receptor expression in the northern house wren (*Troglodytes aedon*), a temperate songbird with female song"

### Table S1

Concentration and purity of RNA extracts, means ± SD. HVC: acronym serves as proper name, RA: robust nucleus of the arcopallium.

| Brain Region | RNA concentration (ng/μL) | Purity ($A_{260}/A_{280}$) |
|---|---|---|
| Area X | 68.35 ± 12.65 | 1.88 ± 0.07 |
| HVC | 94.58 ± 26.55 | 1.93 ± 0.07 |
| RA | 118.12 ± 28.41 | 1.99 ± 0.05 |

### Table S2

Post hoc Tukey's tests comparing differences between brain regions.
AR: androgen receptor, AROM: aromatase, ERα: estrogen receptor α, ERβ: estrogen receptor β.

| Gene | Contrast | Estimate | Std. Error | df | *p*-value | |
|---|---|---|---|---|---|---|
| **AR** | **HVC - Area X** | 2.372 | 0.152 | 77 | **<0.001** | *** |
| | **HVC - RA** | -0.617 | 0.151 | 77 | **<0.001** | *** |
| | **Area X - RA** | 1.754 | 0.152 | 77 | **<0.001** | *** |
| **AROM** | **HVC - Area X** | 6.93 | 0.252 | 77 | **<0.001** | *** |
| | **HVC - RA** | -1.23 | 0.250 | 77 | **<0.001** | *** |
| | **Area X - RA** | 5.70 | 0.252 | 77 | **<0.001** | *** |
| **ERα** | **HVC - Area X** | 3.234 | 0.209 | 77 | **<0.001** | *** |
| | **HVC - RA** | -2.645 | 0.207 | 77 | **<0.001** | *** |
| | **Area X - RA** | 0.589 | 0.209 | 77 | **<0.001** | *** |
| **ERβ** | **HVC - Area X** | 1.44 | 0.122 | 77 | **<0.001** | *** |
| | **HVC - RA** | 0.92 | 0.121 | 77 | **<0.001** | *** |
| | **Area X - RA** | 2.36 | 0.122 | 77 | **<0.001** | *** |

Bolded values are statistically significant based on a *p*-value of 0.05.
Significance codes: '***' $0 \le p \le 0.001$ | '**' $0.001 < p \le 0.01$ | '*' $0.01 < p \le 0.05$ | '.' $0.05 < p \le 0.1$

**Table S3**
Post hoc Tukey's tests comparing differences between breeding females, breeding males, and nonbreeding males.

| Gene | Brain Region | Contrast | Estimate | Std. Error | df | *p*-value | |
|---|---|---|---|---|---|---|---|
| AR | HVC | **Breeding Female – Breeding Male** | 1.260 | 0.230 | 108 | **<0.001** | *** |
| | | Breeding Female – Nonbreeding Male | 0.645 | 0.343 | 108 | 0.150 | |
| | | Breeding Male – Nonbreeding Male | -0.616 | 0.345 | 108 | 0.180 | |
| | Area X | **Breeding Female – Breeding Male** | 0.580 | 0.233 | 108 | **0.038** | * |
| | | Breeding Female – Nonbreeding Male | 0.663 | 0.343 | 108 | 0.134 | |
| | | Breeding Male – Nonbreeding Male | 0.083 | 0.348 | 108 | 0.969 | |
| | RA | **Breeding Female – Breeding Male** | 0.628 | 0.230 | 108 | **0.020** | * |
| | | Breeding Female – Nonbreeding Male | -0.246 | 0.343 | 108 | 0.754 | |
| | | **Breeding Male – Nonbreeding Male** | -0.874 | 0.345 | 108 | **0.034** | * |
| AROM | HVC | Breeding Female – Breeding Male | 0.276 | 0.423 | 102 | 0.792 | |
| | | Breeding Female – Nonbreeding Male | -0.942 | 0.633 | 102 | 0.301 | |
| | | Breeding Male – Nonbreeding Male | -1.218 | 0.637 | 102 | 0.141 | |
| | Area X | Breeding Female – Breeding Male | 0.944 | 0.430 | 103 | 0.7921 | |
| | | Breeding Female – Nonbreeding Male | -0.765 | 0.644 | 102 | 0.301 | |
| | | Breeding Male – Nonbreeding Male | -1.709 | 0.641 | 103 | 0.141 | |
| | RA | Breeding Female – Breeding Male | 0.863 | 0.423 | 102 | 0.109 | |
| | | Breeding Female – Nonbreeding Male | -0.071 | 0.633 | 102 | 0.993 | |
| | | Breeding Male – Nonbreeding Male | -0.933 | 0.637 | 102 | 0.312 | |
| ERα | HVC | **Breeding Female – Breeding Male** | 1.208 | 0.306 | 110 | **<0.001** | *** |
| | | **Breeding Female – Nonbreeding Male** | 1.360 | 0.458 | 110 | **0.010** | * |
| | | Breeding Male – Nonbreeding Male | 0.152 | 0.461 | 110 | 0.942 | |
| | Area X | Breeding Female – Breeding Male | -0.040 | 0.311 | 110 | 0.991 | |
| | | Breeding Female – Nonbreeding Male | -0.037 | 0.458 | 110 | 0.997 | |
| | | Breeding Male – Nonbreeding Male | 0.003 | 0.464 | 110 | 1.000 | |
| | RA | Breeding Female – Breeding Male | 0.429 | 0.306 | 110 | 0.345 | |
| | | Breeding Female – Nonbreeding Male | 0.498 | 0.458 | 110 | 0.523 | |
| | | Breeding Male – Nonbreeding Male | 0.070 | 0.461 | 110 | 0.987 | |
| ERβ | HVC | Breeding Female – Breeding Male | -0.279 | 0.189 | 109 | 0.306 | |
| | | Breeding Female – Nonbreeding Male | -0.271 | 0.283 | 109 | 0.604 | |
| | | Breeding Male – Nonbreeding Male | 0.008 | 0.285 | 109 | 1.000 | |
| | Area X | Breeding Female – Breeding Male | 0.168 | 0.192 | 109 | 0.660 | |
| | | Breeding Female – Nonbreeding Male | -0.020 | 0.283 | 109 | 0.998 | |
| | | Breeding Male – Nonbreeding Male | -0.187 | 0.287 | 109 | 0.792 | |
| | RA | Breeding Female – Breeding Male | -0.258 | 0.189 | 109 | 0.364 | |
| | | Breeding Female – Nonbreeding Male | -0.159 | 0.283 | 109 | 0.841 | |
| | | Breeding Male – Nonbreeding Male | 0.099 | 0.285 | 109 | 0.935 | |

Bolded values are statistically significant based on a *p*-value of 0.05.
Significance codes: '***' $0 \le p \le 0.001$ | '**' $0.001 < p \le 0.01$ | '*' $0.01 < p \le 0.05$ | '.' $0.05 < p \le 0.1$
